# Chinese Glioma Genome Atlas (CGGA): A Comprehensive Resource with Functional Genomic Data for Chinese Glioma Patients

**DOI:** 10.1101/2020.01.20.911982

**Authors:** Zheng Zhao, Ke’nan Zhang, Qiangwei Wang, Guanzhang Li, Fan Zeng, Ying Zhang, Fan Wu, Ruichao Chai, Zheng Wang, Chuanbao Zhang, Wei Zhang, Zhaoshi Bao, Tao Jiang

**Author notes:** Equal contribution (Zheng Zhao, Ke’nan Zhang, Qiangwei Wang). Corresponding authors: (Jiang T), (Bao ZS).

## Abstract

Gliomas are the most common and malignant intracranial tumours in adults. Recent studies have shown that functional genomics greatly aids in the understanding of the pathophysiology and therapy of glioma. However, comprehensive genomic data and analysis platforms are relatively limited. In this study, we developed the Chinese Glioma Genome Atlas (CGGA, http://www.cgga.org.cn), a user-friendly data portal for storage and interactive exploration of multi-dimensional functional genomic data that includes nearly 2,000 primary and recurrent glioma samples from Chinese cohorts. CGGA currently provides access to whole-exome sequencing (286 samples), messenger RNA sequencing (1,018 samples) and microarray (301 samples), DNA methylation microarray (159 samples), and microRNA microarray (198 samples) data, as well as detailed clinical data (e.g., WHO grade, histological type, critical molecular genetic information, age, sex, chemoradiotherapy status and survival data). In addition, we developed an analysis tool to allow users to browse mutational, mRNA/microRNA expression, and DNA methylation profiles and perform survival and correlation analyses of specific glioma subtypes. CGGA greatly reduces the barriers between complex functional genomic data and glioma researchers who seek rapid, intuitive, and high-quality access to data resources and enables researchers to use these immeasurable data sources for biological research and clinical application. Importantly, the free provision of data will allow researchers to quickly generate and provide data to the research community.

## Introduction

Gliomas are the most frequent malignant tumours of the adult brain. According to a multi-centre cross-sectional study on brain tumours in China, the prevalence of primary brain tumours in all populations is approximately 22.52 per 100,000 persons, with gliomas accounting for 31.1% of the population aged 20–59 years [1-3]. According to the histopathological classification of the 2016 World Health Organization (WHO) grading system, glioma is classified from grade II to grade IV by both histological characteristics and several new molecular pathological features, such as *IDH* mutation status and chromosome 1p/19q co-deletion status [4]. Despite advances in current treatment standards, the survival rate of patients with glioma has not changed in decades, especially for aggressive gliomas (with a poor median survival time of only 12 to 14 months) [5, 6]. In addition, most lower-grade gliomas (grade II and III, LGG) will progress to glioblastoma (grade IV, GBM) in less than 10 years [4, 7, 8]. At present, the reasons for glioma recurrence or malignant progression may be as follows: 1) infiltrative tumour cells cannot be completely removed by neurosurgical resection [9, 10]; 2) retained tumour cells cannot be effectively suppressed by limited postoperative treatment options [3, 11, 12]; 3) multiple lesions may develop [13, 14]; 4) cell cloning is rapid under chemotherapy and/or radiotherapy [7, 15]; 5) the adaptive tumour microenvironment permits tumour cells [16, 17]; and 6) limited data resources lead to limited research. Therefore, it is essential to collect clinical specimens and generate genomic data for the glioma research community.

Recent high-throughput technologies have enabled extensive characterization of genomic status, including but not limited to DNA methylation modification, genetic alteration, and gene expression regulation. In the cancer research community, major large-scale projects, such as The Cancer Genome Atlas (TCGA, including 516 LGGs and 617 GBMs before Oct. 18, 2019) [18] and the International Cancer Genome Consortium (ICGC, excluding TCGA samples, including 80 adult GBMs and 50 paediatric GBMs before April. 3, 2019) [19, 20], have generated an unparalleled amount of functional genomic data. These projects have begun to transform our understanding of cancer and even lead to improvements in our ability to diagnose, treat, and prevent human cancers. Importantly, they have provided an opportunity to make and validate important discoveries for cancer genomic researchers around the globe. However, the data resources generated by these projects are often not easy to access directly, analyse or visualize, especially for researchers with no bioinformatics skills, thus preventing the translation of functional genomics results into novel findings of biological significance for drug development and clinical treatment. Although several webservers, such as cBioportal [21, 22] and GlioVis [23], have been built to integrate analysed multi-dimensional glioma data, they have ignored the presence of cancer heterogeneity in gliomas, which cannot be examined in specific subtypes and is rarely found in recurrent glioma samples.

Here, we introduce the CGGA (Chinese Glioma Genome Atlas, http://www.cgga.org.cn) database, which is an open-access and easy-to-use platform for interactive exploration of multi-dimensional functional genomic datasets for nearly 2,000 primary and recurrent glioma samples from Chinese cohorts. CGGA currently contains whole-exome sequencing (286 samples), messenger RNA (mRNA) sequencing (1,018 samples), microarray (301 samples), DNA methylation microarray (159 samples), microRNA microarray (198 samples) and comprehensive clinical data. We also developed an analysis module to allow users to browse the mutational landscape profile, mRNA/microRNA expression profile and DNA methylation profile as well as to perform survival and correlation analyses for specific glioma subtypes. We believe that this website will greatly reduce the barriers between complex functional genomic data and glioma researchers who seek rapid, intuitive, and high-quality access to data resources.

## Results

### Database content and usage

The CGGA database was designed to store functional genomic data and to allow interactive exploration of multi-dimensional datasets from primary and recurrent gliomas in Chinese cohorts; it is available at http://www.cgga.org.cn/. Currently, CGGA contains whole-exome sequencing data (286 samples), messenger RNA sequencing data (total: 1,018 samples, batch 1 with 693 samples and batch 2 with 325 samples), microarray data (301 samples), DNA methylation microarray data (159 samples), and microRNA microarray data (198 samples) for glioma. The database also contains detailed clinical data (including WHO grade and histological type, critical molecular genetic information, age, sex, chemoradiotherapy status and survival data). Detailed statistical information for each dataset is provided in Table 1. We organized the web interface of CGGA according to the three main functional features: (i) Home, (ii) Analyse, and (iii) Download. In the following context, we provide an example for using CGGA.

**Table 1.**
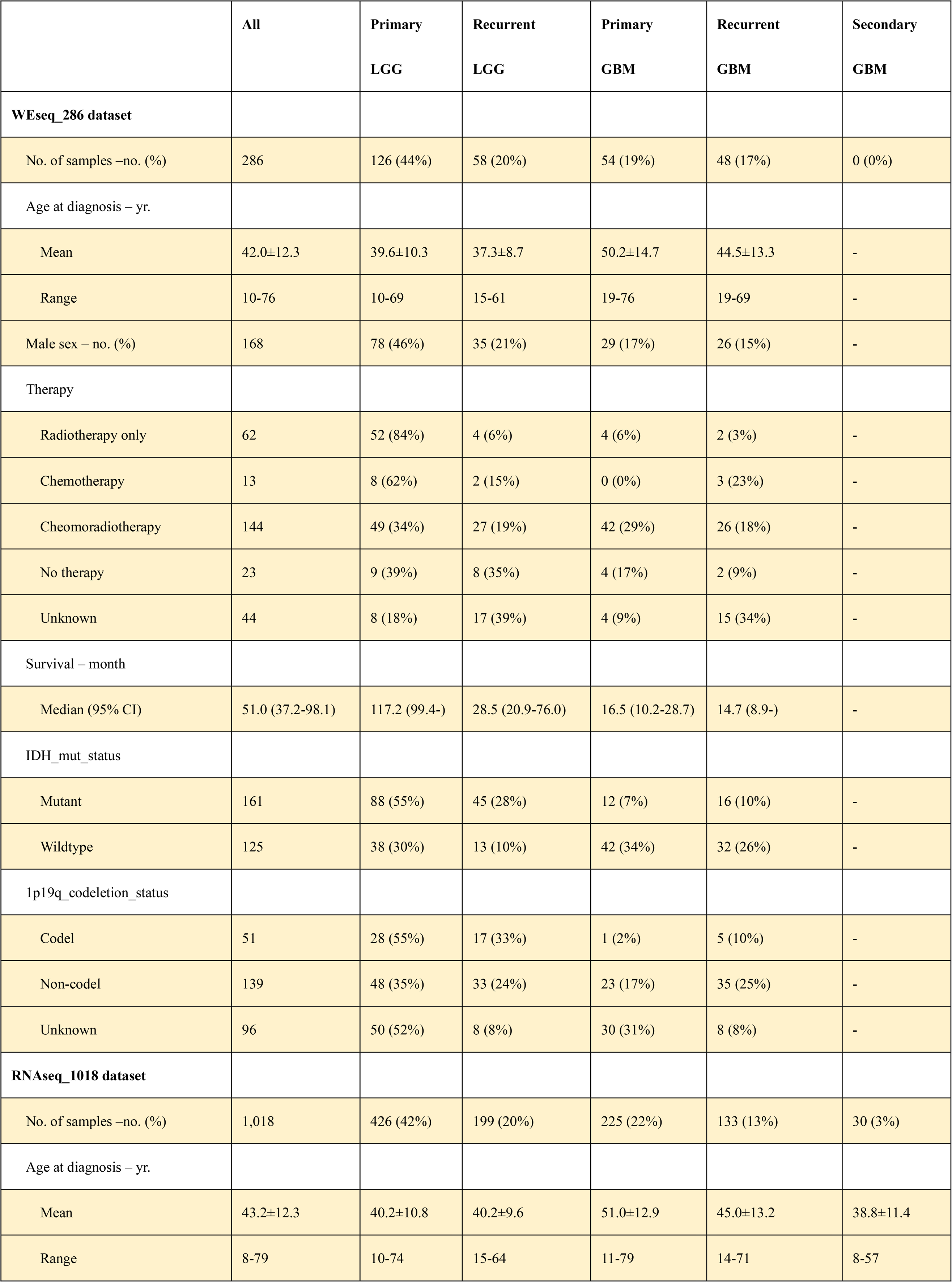
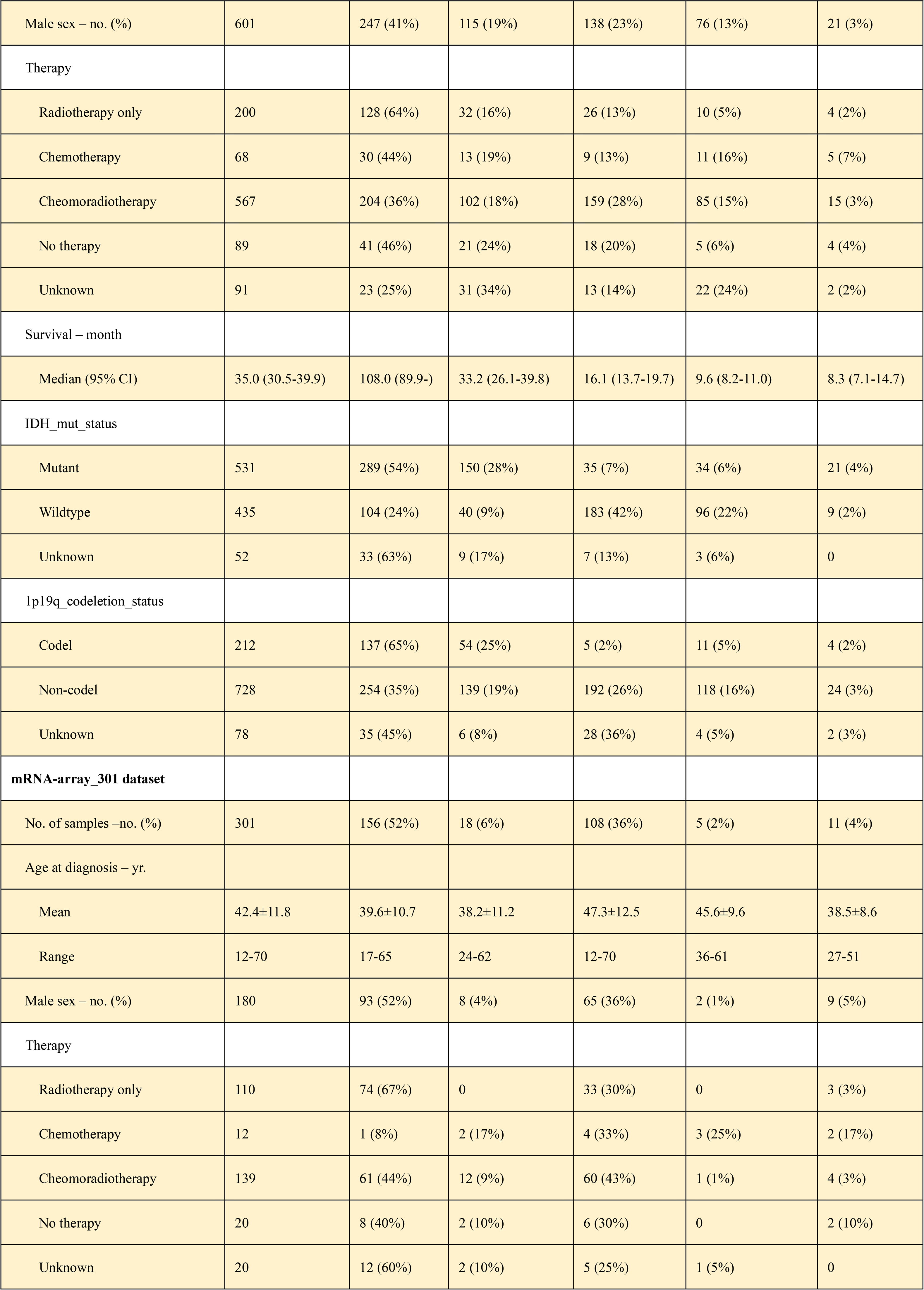
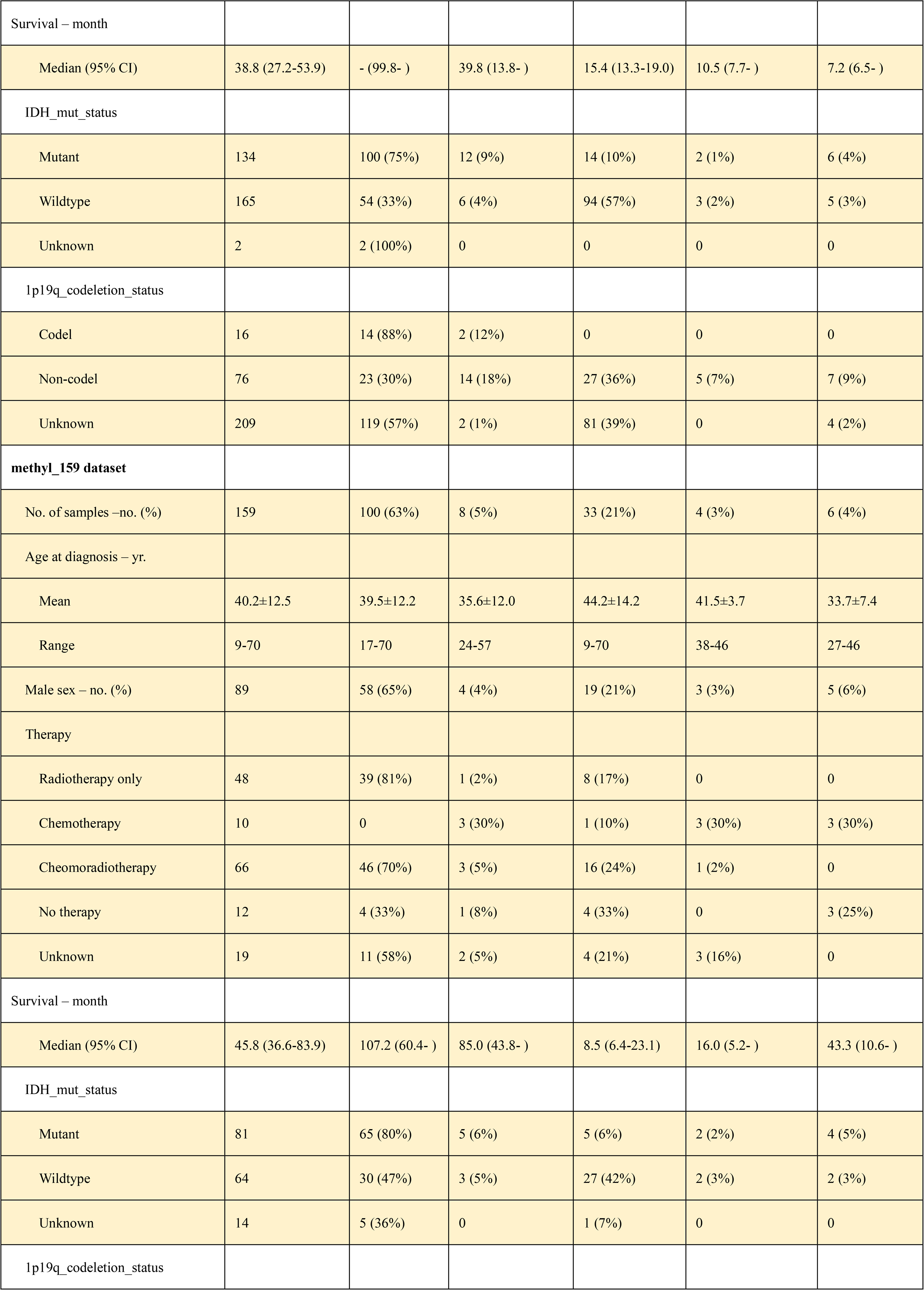
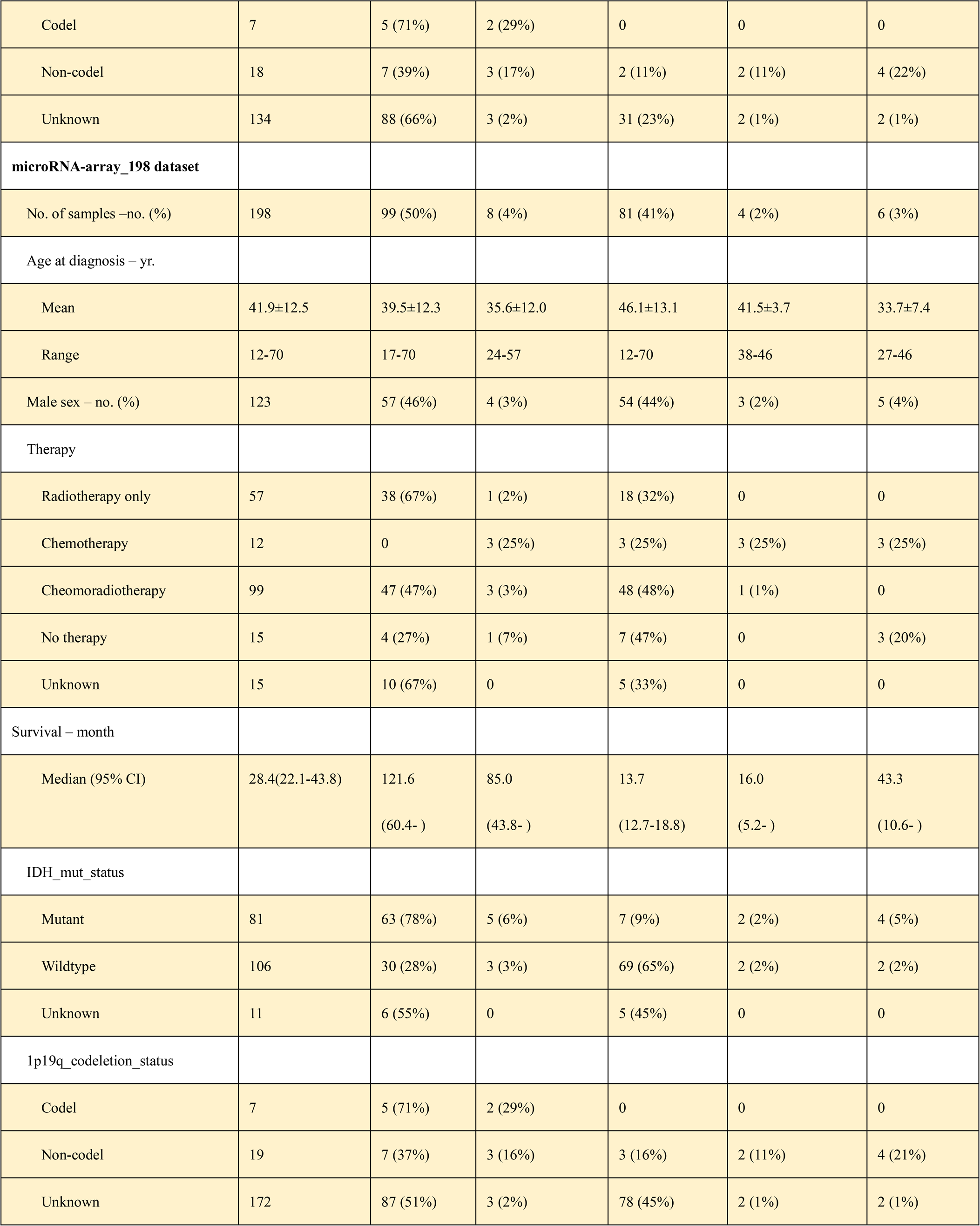
Clinical and Phenotypical Characteristics of Data Set in CGGA database

### The home page

On the ‘Home’ page, CGGA provides a statistical table for a glioma dataset, including the dataset name, data type, number of samples in each subgroup, clinical data and analysis purposes. For instance, we performed messenger RNA sequencing on 1,018 glioma samples included in two datasets (693 samples in batch 1 and 325 samples for batch 2, including 282 primary LGGs, 161 recurrent LGGs, 140 primary GBMs and 109 recurrent GBMs in batch 1 and 144 primary LGGs, 38 recurrent LGGs, 85 primary GBMs, 24 recurrent GBMs and 30 secondary GBMs in batch 2). To the best of our knowledge, CGGA is the first database to store the functional genomic data for both LGG and GBM recurrent gliomas. In addition, users can obtain a visualized result for the analysis of each dataset for a specific glioma subtype by clicking on a hyperlink on the ‘Home’ page. The ‘Download’ and ‘Help’ pages can also be accessed directly from the ‘Home’ page.

### Overall analyses and results

To facilitate analysis of the CGGA data by researchers, we developed four online modules in the ‘Analyse’ tab, including ‘WEseq data’, ‘mRNA data’, ‘methylation data’, and ‘microRNA data’, to analyse whole-exome, mRNA expression, DNA methylation and microRNA expression data, respectively (Figure 1A). A key feature of CGGA is that it is easy to use. In the context below, we demonstrate the use of the ‘Analyse’ tab in CGGA.

**Figure 1.**
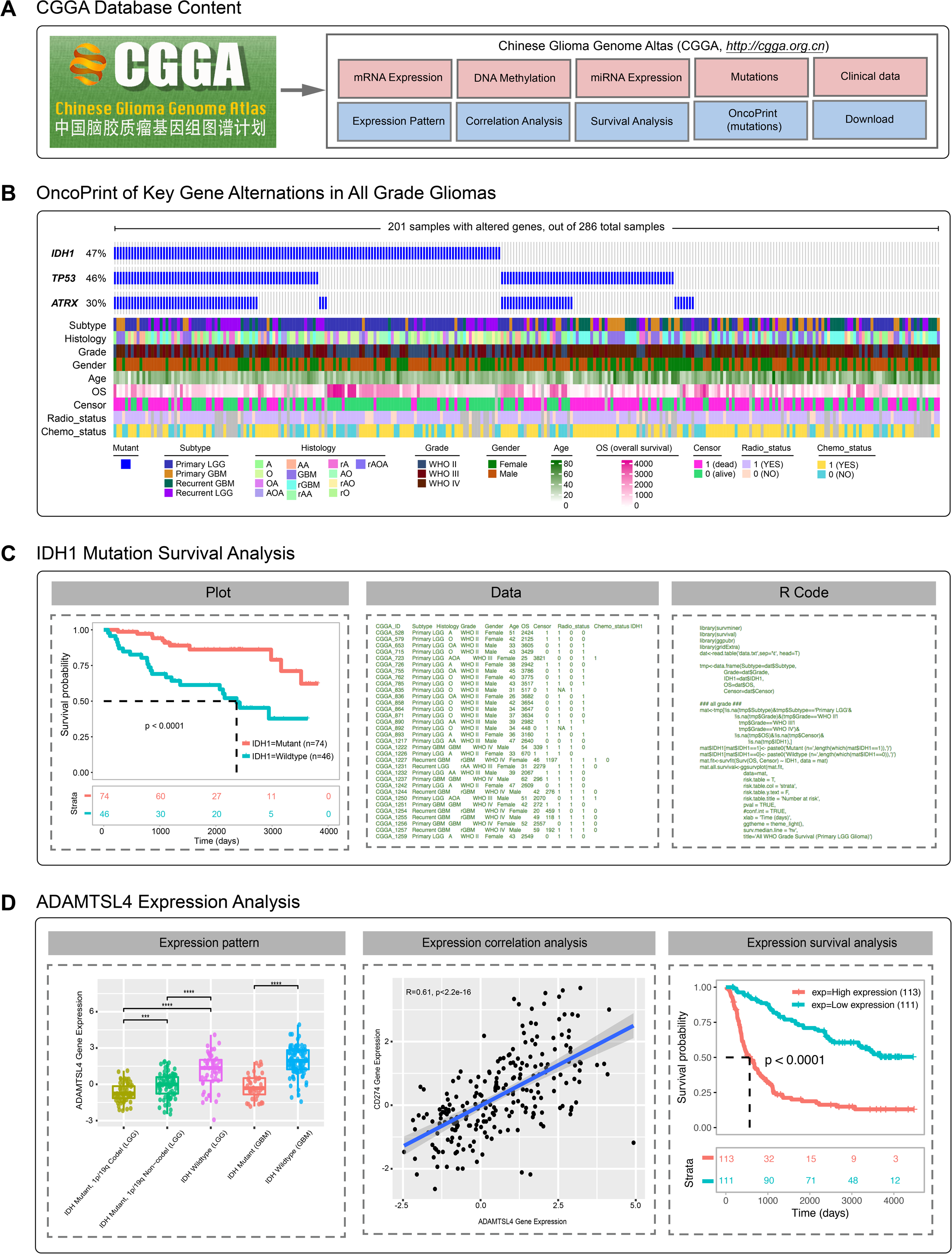
An overview of the CGGA database. A. The CGGA contains whole-exome sequencing, mRNA and microRNA expression, and DNA methylation data, clinical data, and several analysis modules; B. The mutation profile in all gliomas (in the ‘WSseq_286’ dataset); C. left: the overall survival of glioma patients with IDH1 mutation and the wild-type gene from primary LGGs (in the ‘WSseq_286’ dataset); middle: the data was used to generate the plot; right: the R code was used to generate the plot; D. left: the ADAMTSL4 gene expression distribution in primary gliomas based on 2016 WHO grading system (in the ‘mRNAseq_325’ dataset); middle: the gene expression correlation between ADAMTSL4 and CD274 genes (using ‘mRNAseq_325’ dataset); right: the overall survival of glioma patients with low and high ADAMTSL4 gene expression (in the ‘mRNAseq_325’ dataset).

On the ‘WEseq data’ page, users are allowed to visualize the mutational profile of a gene set of interest and survival analysis of a specific gene of interest in a specific glioma subtype. In the ‘Oncoprint’ section, users are guided to a) input a gene set of interest (‘*IDH1 TP53 ATRX’* for example), and b) select a dataset of interest (‘All’ for example). Based on user input, this tool automatically generates visualized results. In this result, each case or patient is represented as columns, each gene is displayed as rows, and a colour map on the bottom is used to depict specific clinical information (Figure 1B). This ‘Oncoprint’ can be very useful for visualizing the mutational profile for a gene set of interest in a specific glioma subtype and for intuitively validating trends such as mutational frequency and mutual exclusivity or co-occurrence for a gene pair. In the above example, mutations in the *IDH1* (47%), *TP53* (46%) and *ATRX* (30%) genes were the most common mutations in all gliomas. In the ‘Survival’ section, users are allowed to a) input a specific gene of interest (‘*IDH1’* for example), and b) select a dataset of interest (‘Primary LGG*’* for example) to investigate the association of the mutation with severe functional consequences. Consistent with previous studies [24], primary LGG cases with *IDH1* mutations have a better overall survival than do cases with *IDH1* wild-type tumours (p < 0.0001, Figure 1C, left). These analysis results from the ‘WEseq data’ section can be exported as a PDF file. For the sake of reproducibility, we provide the analysis data (Figure 1C, middle) and R code (Figure 1C, right), which allow users to reproduce the figure to be able to modify or adapt each figure according to each researcher’s demands.

On the ‘mRNA data’ page, users are allowed to perform gene expression distribution, correlation and survival analyses for a specific gene of interest in a specific glioma subtype. Three mRNA datasets are available for users, including two batch RNA-seq datasets (batch 1: 693 samples; batch 2: 325 samples) and one microarray dataset (301 samples). In the ‘Distribution’ section, users can display one gene distribution pattern for each glioma subtype by selecting a dataset (‘mRNAseq_325’ for example) and inputting a gene name of interest (‘*ADAMTSL4’* for example). The results show the gene expression pattern in each glioma subtype classified by clinical information. Similar to our previous studies [25], the *ADAMTSL4* gene was shown to be differentially expressed according to the WHO 2016 classification based on the *IDH* mutation and/or 1p/19q co-deletion status (Figure 1D, left). Moreover, a critical feature of the CGGA dataset is the inclusion of recurrent gliomas. This module allows users to infer whether a gene may be a candidate factor that drives malignant progression if it is differentially expressed in primary and recurrent gliomas. In the ‘Correlation’ section, the user is allowed to validate the co-expression pattern by selecting a dataset (‘mRNAseq_325’ for example) and inputting a gene pair (‘*ADAMTSL4*’ and ‘*CD274*’ for example). As a result, the co-expression patterns in each glioma subtype will be displayed with the results of Pearson’s test and the p value (Figure 1D, middle). In the ‘Survival’ section, users can perform survival analysis based on gene expression by selecting a dataset (‘mRNAseq_325’ for example) and inputting a gene of interest (‘*ADAMTSL4*’ for example). All primary glioma patients with low *ADAMTSL4* expression showed better overall survival than did those with high *ADAMTSL4* expression (p < 0.0001, Figure 1D, right). The above results from the ‘mRNA data’ section are consistent with our previous study [25]. Similar to the ‘mRNA data’ page, users can also display the methylation/microRNA distribution and perform correlation and survival analyses on the ‘methylation data’ page and the ‘microRNA data’ page, respectively.

### Data acquisition

All the data sets in CGGA can be downloaded on the ‘Download’ page by both the community and researchers. Each data type is saved at the gene and/or probe level and is then combined with the available clinical data, including basic clinical information, survival and therapy information.

### Perspectives and concluding remarks

The current version of the CGGA is the first release of our database, and it incorporates multi-dimensional functional genomic glioma data, including whole-exome sequencing, mRNA and microRNA expression, and DNA methylation data for nearly 2,000 samples from Chinese cohorts. Considering the importance of these data for glioma research, CGGA is publicly available. To the best of our knowledge, CGGA is the first database to store the functional genomic data for both recurrent LGGs and GBMs. In addition, CGGA provides several tools that allow users to analyse these datasets, including mutational profile, distribution pattern, correlation and survival analysis tools. These tools will be useful for users to generate or validate findings of novel biological significance.

We anticipate several future directions for our CGGA database. First, through the Beijing Neurosurgical Institute, Beijing Tiantan Hospital and Chinese Glioma Cooperative Group (CGCG) Research Network, we will continue to collect glioma samples and perform multiple ‘Omics’ sequencing/microarray analyses, and we will continue to update this database regularly in the future. Second, we also plan to add image-genomic data that match the ‘Omics’ data in CGGA. Third, we will develop more advanced features, including data for other ‘Omics’ analyses, search functions for clinical information on a patient of interest, and further extensions for the data analysis tools. In summary, CGGA facilitates access to functional genomic data for Chinese cohorts for the entire glioma community. It provides an easy-to-use, user-friendly interface for obtaining integrated data sets, performing intuitive visualized analysis, and downloading these datasets. CGGA greatly reduces the barriers between complex functional genomic data and glioma researchers, which empowers researchers to use functional genomic data into important biological insights and potential clinical applications.

## Materials and methods

### Clinical specimen collection

Glioma tissues, corresponding genomic data and patient follow-up information were obtained from Beijing Tiantan Hospital at Capital Medical University, Tianjin Medical University General Hospital, Sanbo Brain Hospital at Capital Medical University, the Second Affiliated Hospital of Harbin Medical University, the First Affiliated Hospital of Nanjing Medical University, and the First Hospital of China Medical University. All research performed was approved by the Beijing Tiantan Hospital Capital Medical University Institutional Review Board (IRB) and was conducted according to the principles of the Helsinki Declaration. According to the central pathology reviews of independent committee certified neuropathologists, all the subjects were consistently diagnosed with glioma and further classified according to the 2007/2016 WHO classification system. All patients provided written informed consent. The specimens were collected under IRB KY2013-017-01 and frozen in liquid nitrogen within 5 min of resection.

### Data processing for whole-exome sequencing data

Genomic DNA from tumours and the matched blood samples was extracted, and high integrity was confirmed by 1% agarose gel electrophoresis. The DNA was subsequently fragmented and quality-controlled, and paired-end libraries were prepared. Agilent SureSelect kit v5.4 was used for target capture. Sequencing was performed using the Illumina HiSeq 4000 platform with a paired-end sequencing strategy. Valid DNA sequencing data were mapped to the reference human genome (UCSC hg19) using Burrows-Wheeler Aligner (v0.7.12-r1039, bwa mem) [26] with default parameters. SAMtools (version 1.2) [27] and Picard (version 2.0.1, Broad Institute) were then used to sort the reads by coordinates and mark duplicates. Statistics such as sequencing depth and coverage were calculated based on the resulting BAM files. SAVI2 was used to identify somatic mutations (including single-nucleotide variations and short insertions/deletions) as previously described [7, 8]. Briefly, in this pipeline, SAMtools mpileup and bcftools (version 0.1.19) [28] were employed to perform variant calling, and the preliminary variant list was filtered to remove positions with no sufficient sequencing depth, positions with only low-quality reads, and positions biased toward either strand. Somatic mutations were identified and evaluated by an empirical Bayesian method. In particular, mutations with a significantly higher mutation allele frequency in tumours than in normal controls were selected.

### Data processing for mRNA sequencing data

Prior to library preparation, total RNA was isolated using RNeasy Mini Kit (Qiagen) according to the manufacturer’s instructions. A pestle and QIAshredder (Qiagen) were used to disrupt and homogenize frozen tissue. The RNA intensity was checked using a 2,100 Bioanalyzer (Agilent Technologies), and only high-quality samples with an RNA integrity number (RIN) value greater than or equal to 6.8 were used to construct the sequencing library. Typically, 1 μg of total RNA was used with the TruSeq RNA library preparation kit (Illumina) in accordance with the low-throughput protocol, except that SuperScript III reverse transcriptase (Invitrogen) was used to synthesize first-strand cDNA. After PCR enrichment and purification of adapter-ligated fragments, the concentration of DNA with adapters was determined by quantitative PCR (Applied Biosystems 7,500) using primers QP1 5’-AATGATACGGCGACCACCGA-3’ and QP2 5’-CAAGCAGAAGACGGCATACGAGA-3’. The length of the DNA fragment was measured using a 2,100 Bioanalyzer, with median insert sizes of 200 nucleotides. The RNA-seq libraries were sequenced using the Illumina HiSeq 2,000, 2,500 or 4,000 Sequencing System. The libraries were prepared using the paired-end strategy with read lengths of 101 bp, 125 bp or 150 bp. Base calling was performed by the Illumina CASAVA v1.8.2 pipeline. RNA-seq mapping and quantification were processed by using STAR (version v2.5.2b) [29] and RSEM (version 1.2.31) software [30]. Briefly, reads were aligned to the human genome reference (GENCODE v19, hg19) with STAR, and then sequencing read counts for each GENCODE gene were calculated using RSEM. The expression levels of different samples were merged into an FPKM (fragments per kilobase transcriptome per million fragments) matrix. We defined a gene as expressed only if its expression level was greater than 0 in half of the samples. Finally, we retained only expressed genes in the mRNA expression profile.

### Data processing for mRNA microarray data

A rapid haematoxylin & eosin stain for frozen sections was performed on each sample to assess the tumour cell proportion before RNA extraction. RNA was extracted from only samples with >80% tumour cells. Total RNA was extracted from frozen tumour tissue with the mirVana miRNA Isolation Kit (Ambion), as described previously [31]. A NanoDrop ND-1000 spectrophotometer (NanoDrop Technologies) was used to evaluate the quality and concentration of extracted total RNA and an Agilent 2100 Bioanalyzer (Agilent) to assess the integrity. The qualified RNA was collected for further processing. cDNA and biotinylated cRNA were synthesized and hybridized to Agilent Whole Human Genome Array according to the manufacturer’s instructions. Finally, the array-generated data were analyzed by the Agilent G2565BA Microarray Scanner System and Agilent Feature Extraction Software (Version 9.1). GeneSpring GX11.0 was applied to calculate the probe intensity.

### Data processing for methylation microarray data

A haematoxylin and eosin-stained frozen section was prepared for assessment of the percentage of tumour cells before RNA extraction. Only samples with greater than 80% tumour cells were selected. Genomic DNA was isolated from frozen tumour tissues using the QIAamp DNA Mini Kit (Qiagen) according to the manufacturer’s protocol. The DNA concentration and quality were assessed using a NanoDrop ND-1000 spectrophotometer (NanoDrop Technologies, Houston, TX). The microarray analysis was performed using Illumina Infinium HumanMethylation27 Bead-Chips (Illumina Inc.), which contains 27,578 highly informative CpG sites covering more than 14,000 human RefSeq genes. This allows researchers to investigate all sites per sample at a single-nucleotide resolution. Bisulfite modification of DNA, chip processing and data analysis were performed following the manufacturer’s manual at Wellcome Trust Centre for Human Genetics Genomics Lab, Oxford, UK. The array results were examined with the BeadStudio software (Illumina).

### Data processing for microRNA microarray data

Total RNA (tRNA) was extracted from frozen tissues by using the mirVana miRNA Isolation Kit (Ambion, Inc., Austin, Tex), and the concentration and quality were determined with a NanoDrop ND-1000 spectrophotometer (NanoDrop Technologies, Wilmington, Del). microRNA expression profiling was performed using the human v2.0 microRNA Expression BeadChip (Illumina, Inc., San Diego, Calif) with 1146 microRNAs covering 97% of the miRBase 12.0 database according to the manufacturer’s instructions.

### Implementation

In CGGA, all data are organized with MySQL 14.14 based on relational schema, which will be supported by future CGGA updates. The website code was written based on Java Server Pages using the Java Servlet framework. The website is deployed on the Tomcat 6.0.44 web server and runs on a CentOS 5.5 Linux system. JQuery was used to generate, render and manipulate data visualization. The ‘Analyse’ module was realized with Perl and R scripts. The CGGA website has been fully tested in Google Chrome and Safari browsers.

## Data Availability

All data for this article can be found online at http://www.cgga.org.cn.

## Authors’ contributions

TJ, ZB, WZ, and ZZ conceived and supervised this study. ZZ, KZ and QW designed the research. ZZ, GL, FZ, YZ, FW, RC, ZW, CZ performed data analysis. ZZ developed CGGA web server. TJ, ZB, and ZZ wrote the manuscript. All authors read and approved the final manuscript.

## Acknowledgments

This work was supported by the National Natural Science Foundation of China (NSFC) fund (Nos. 81702460 and 81802994).

